# Intrinsic noise reveals the stability of a neuronal network

**DOI:** 10.1101/2025.07.23.666219

**Authors:** Marcelo Bussotti Reyes, Ramon Huerta, Pedro Valadão Carelli, Reynaldo D. Pinto, Mikhail I. Rabinovich, Allen I. Selverston

## Abstract

The stability of rhythmic activity in neural networks is an important aspect in the study of central pattern generators (CPGs). Different from other physiological rhythms, the activity of CPGs has not been fully characterized in terms of its stability, especially using quantitative methods. We propose a method that takes advantage of the natural noise present in CPGs to quantify the stability of the rhythmic activity. Furthermore, we used the stationary bootstrap method to define confidence intervals of the results. We applied this method to study the influence of a synaptic modification on the pyloric CPG circuit, using artificial synapses implemented in dynamic clamp software. We show that even after removing one of its strongest synapses, the CPG stability remains unaltered. This analysis suggests that CPGs are designed to be strongly stable regardless of the parameter perturbations they undergo.

## Introduction

The stability of the rhythmic activity of neuronal networks is a central issue in the study of central pattern generators (CPGs). These networks possess the remarkable capability of producing robust behavior, maintaining their activity under different kinds of perturbations such as modulation from higher centers (Pflüger, 1999; Dickinson, 2006), sensory feedback (Rossignol et al., 2006), noise introduced by random fluctuations of ionic channel states (Carelli et al., 2005), or even with respect to strong synaptic modifications (Reyes et al., 2008). The source of such extraordinary robustness observed in CPGs is still not understood and has interesting applications to the understanding of more complex networks and for applied fields such as robotics (Fukuoka et al., 2003; Gutierrez-Galan et al., 2020).

A lot of effort has been made to quantitatively analyze physiological systems, as for example heartbeat and blood pressure. Much of this analysis turned out to be successful in discriminating healthy from ill individuals exclusively from dynamical or statistical properties of the time series. Similar methods were also able to improve the predictability of sudden heart attacks (Herd, 1991; Camm et al., 1996; Kudaiberdieva et al., 2007; Glass, 2001). However, not the same effort has been made to develop methods for understanding the dynamics of CPGs.

In simple CPGs, it is possible to understand the role of individual neurons and synapses in the stability of the rhythmic pattern (Weaver and Hooper, 2003; Mamiya and Nadim, 2004). In more complex networks, where one can assess only a small portion of the neurons, the search for the mechanisms underlying the pattern generation is mostly made through simulations (Smith et al., 2000; Rybak et al., 2006; Reyes et al., 2015). However, a more complete understanding of the pattern stability remains still in a qualitative fashion.

In contrast to previous work, here we turned our attention to a quantitative analysis of CPGs activity. We used the endogenous noise present in the activity of these networks to estimate the stability of time series. We wanted to quantify the stability of the system not by looking at the small jitter of the periodic oscillations of a pyloric neuron, but by analyzing the stability from the dynamical systems perspective (Escot and Sandubete, 2023; Oku and Aihara, 2018).

The fact that we are dealing with noise requires a probability of ascertain the result. It means that the result of the stability is not completely significant *per se*, but it requires an estimation of the confidence interval. We used the stationary bootstrap method to create a statistical test that calculates the confidence interval of the results. The method also estimates the level of significance of the network stability. Similar approaches used VAR model with the stationary bootstrap (Beyaztas and Abdel-Salam, 2022) in unemployment and inflation data, and used the maximum eigenvalue of the variance-covariance matrix of the multivariate time series to identify regime shift in ecosystems (Brock and Carpenter, 2006).

Finally, we addressed how a synaptic modification impacted the network robustness. We applied a method that analyzed the influence of modifying the synapse from the lateral pyloric (LP) to the pyloric dilator (PD) neuron. This synapse has been shown to play an important role for the stabilization of the pyloric rhythm (Weaver and Hooper, 2003; Mamiya and Nadim, 2004). This question is directly connected to the understanding of how individual components of the network contribute to its final output.

Even though we applied the method to the single case of the pyloric CPG, the method can be extended to other systems, provided that they present spontaneous or sustained stereotyped activity and that one can measure important discrete variables of the system from the time series.

## Materials and Methods

### In Vitro Experiments

The data shown here were already used for a different analysis and published elsewhere (Reyes et al., 2008). We provide a short description of the methods, enough for understanding the results. Further details about the experiments can be found in the previous publication.

Overall, 11 adult California spiny lobsters were used in the experiment. The LP and two PD neurons were impaled with sharp intracellular microelectrodes (15–25 MΩ) filled with either 4 M potassium acetate with 0.1 M KCl or 3 M KCl. One of the PDs was impaled with an extra electrode for current injection (Fig. 1). A modified version of the Dynamic Clamp developed by Pinto et al. Pinto et al. (2001) was used to manipulate the synapse from the LP to PD neuron.

**Figure 1:**
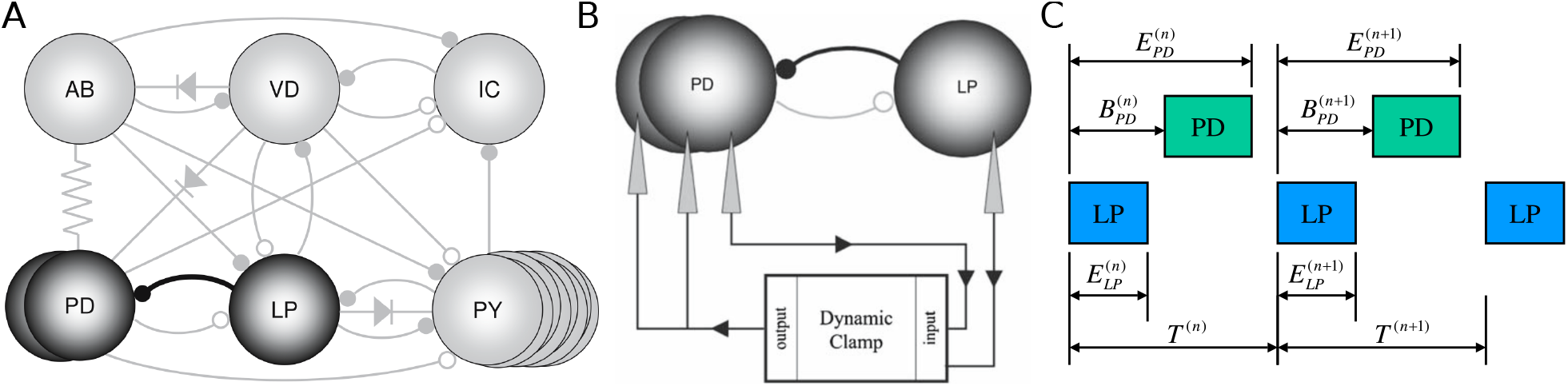
A) Schematic of the crustacean pyloric CPG showing the neurons and the synaptic connections between them. B) Schematic of the dynamic clamp configuration used in the experiments. The LP neuron and one PD neuron were recorded. The Dynamic Clamp system uses the membrane potential recordings from these two neurons to compute the synaptic current, which is then injected into both PD neurons. C) Definition of the variables used in the experiment, illustrated over two oscillation cycles. Each square represents a neuronal burst, measured from the first to the last spike. The variables 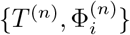 include the cycle period *T*^(*n*)^ in the cycle *n*, defined using the first spike of the LP neuron as a reference, as well as the beginning *B* and the end *E* of the bursts of the LP and PD neurons. Accordingly, the complete set of variables is 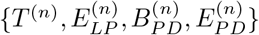

The artificial synaptic current was injected with opposite polarity to the natural inhibitory LP→PD synapse. Increasing values of |*g*_*s*_*yn*| therefore correspond to progressive cancellation and eventual reversal of the native synaptic current. For clarity, all reported values refer to the absolute magnitude of *g*_*syn*_.

The synaptic strength of the LP to PD neuron synapse was modified by choosing the synaptic parameters so that the Dynamic Clamp injected a current that modifies the natural synaptic current. Depending on the artificial synaptic conductance (*g*_*syn*_) the original postsynaptic current could be canceled or even reversed. The synaptic current injected by the Dynamic Clamp was calculated as Destexhe et al. (1994):

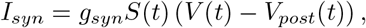

where

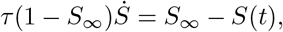

and

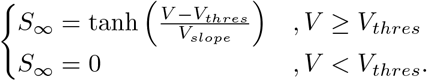

The synaptic threshold for transmitter release *V*_*thres*_ was chosen to match the average of the LP neuron’s membrane potential. The equation parameters were *τ* = 10*ms, V*_*sl*_ = 10*mV*, and *V*_*syn*_ = −80*mV*. Figure 2a shows one example of control time series and one of the modified system, after turning on the dynamic clamp. The synaptic conductance *g*_*syn*_ was modified from 0 (control) to 100nS, and the number of intermediate points in the range varied in different experiments. We referred only to the absolute value of *g*_*syn*_, which ranged from 0 to 100nS. The data were collected from a total of 11 experiments.

**Figure 2:**
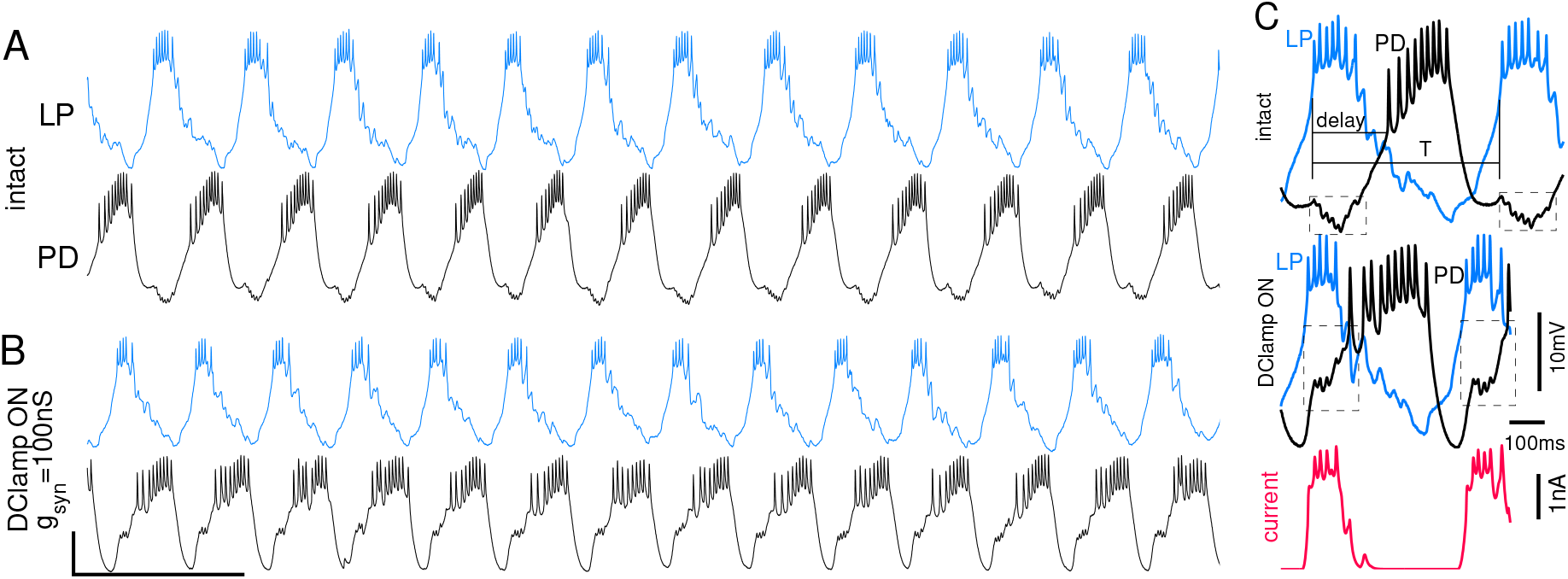
Time series of the recorded membrane potentials of the neurons and definition of the dynamical variables. A) Membrane potential of the LP (upper) and PD (lower trace) neurons in the intact network. B) Same recordings when the dynamic clamp is turned on with *g*_*syn*_ = 100*nS*. Amplification of two consecutive bursts of the LP neuron for the intact rhythm (top) and when the dynamic clamp is turned on. The bottom trace shows the current calculated by the dynamic clamp and injected in both PD neurons. Adapted from Reyes et al. (2008).

We eliminated cycles where one of the neurons did not fire and bursts during or adjacent (6 s before and after) to bursts of the oesophageal neurons (visually identified). After these bursts were eliminated, we calculated the Euclidean distance between two consecutive bursts Σ_*i*_ (*δ*_*i*+1_ − *δ*_*i*_)^2^ and eliminated the measurements that were more distant than 3 SD from the mean. This procedure was used to filter the more local dynamics and eliminate spurious points. When the remaining number of consecutive bursts was smaller than 40, the time series were not used for the analysis.

The beginning and the end of each neuron burst were detected by the first and last action potential in that burst, respectively. The first action potential of the LP neuron was used as the reference for the beginning of the cycle (see Figure 2b).

### Stability Analysis

The neural data we are dealing clearly show two very distinct timescales, one for the spikes and the other for the bursts. Since we are dealing with motoneurons, the phases in which the neurons fire are of particular importance, because they determine the phase of muscle activation. Hence, instead of working with the continuous system, we can analyze the stability of the system described in terms of cycle-related discrete variables. A natural set of these variables are the cycle durations *T*^*n*^ and the phase lags 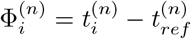, and *i* is the index of all the neurons and phases that can be measured (see Figure 1). The time 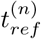 represents the instant of the first spike of the *n*_*th*_ cycle of the neuron considered as the reference. Therefore, the whole set of variables needed to describe the state of the system at the cycle *n* is

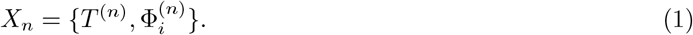

In our experiments, we used simultaneous recordings of the LP and one of the PD neurons for data analysis. The two PD neurons activate synchronously due to strong gap junction connections; therefore, we used the phase of the single PD as the same for both neurons and for the calculation of the synaptic current. We considered the LP as the reference neuron, and all the other variables were measured with respect to its first spike in each cycle. The time of the last spike of the LP (*E*_*LP*_) was used as Φ_1_, the first spike of the PD neuron (*B*_*PD*_)was Φ_2_ and the last spike of the PD (*E*_*PD*_) was Φ_3_.

Once we have the set of discrete variables, we can define a map of the form *X*_*n*+1_ = **F**(*X*_*n*_), where **F** is in general a nonlinear function. The map represents the idea that the state of the system in the *n*_*th*+1_ cycle can be predicted by the state in the previous (*n*_*th*_) cycle, as done in auto-regressive models.

Since the CPG dynamics are close to periodic behavior, the description of the whole dynamics could be reduced to a linear map around the fixed point *X*\*, i.e *δ*_*n*+1_ = **J***δ*_*n*_, where **J** is the Jacobian of the system calculated in the fixed point and *δ*_*n*_ = *X*_*n*_ − *X*\*.

The estimation of the Jacobian matrix was done by taking the dynamical variables *X*_*n*_, subtracting their mean to obtain *δ*_*n*_ and building two separate non-square matrices of same sizes, one containing the variables in the cycle *n* and the other containing the iterations of these variables. Hence, we have two new sets of variables, *Z* = {*δ*_*n*_} and *W* = {*δ*_*n*+1_}. We calculated the parameters [*J*_*ij*_] that best fit the equation by solving the equation *W* = **J***Z*, and we estimated its value as *J*_*est*_ = (*Z*^*T*^ *Z*)^−1^*Z*^−1^*W*, where ^*T*^ stands for the transpose and ^−1^ for the inverse of the matrices.

We used the Stationary Bootstrap algorithm Politis and Romano (1994) to obtain a smooth distribution of eigenvalues and to define a confidence level for the stability analysis of the system.

We re-sampled the series {*X*_*n*_} and estimated the eigenvectors from each of these samples in order to obtain their distributions. The bootstrap method was repeated 10000 times. The distribution of the highest absolute real eigenvalue *max* {*Re*(*λ*_*i*_)} was used to determine the mean and the confidence intervals (Fig. 3).

**Figure 3:**
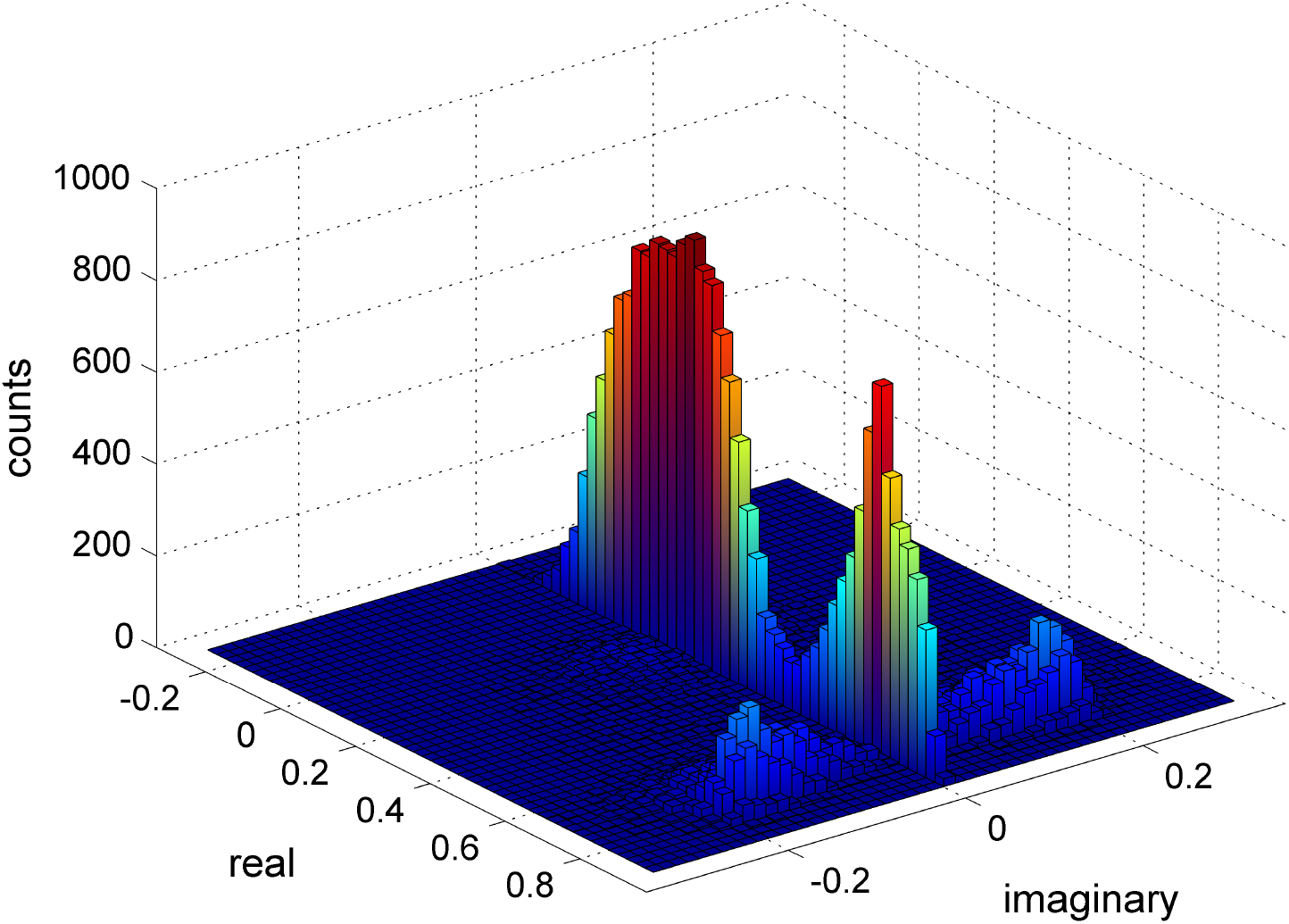
Two-dimensional histogram of the eigenvalues. We re-sampled a series of 996 bursts using the 10000 stationary bootstrap method. Such method produces a smooth distribution of the values, allowing for an estimation of the confidence interval of the measurements. The two higher peaks along the *im*(*λ*) = 0 axis represent the real eigenvalues and the symmetrically positioned show the two complex ones.

## Results

### Application to the pyloric CPG circuit

We have employed our algorithm to study the stability of the pyloric CPG. We characterized the system using the membrane potential time series of the LP and PD neurons. We assume that the whole pyloric network is a dynamical system whose stability can be inferred from the time series from these two neurons. This is reasonable because the PD and LP neurons play important roles in the pyloric CPG: the PDs are part of the pacemaker group, responsible for the main rhythmic activity (around 1 Hz), while the LP projects the only chemical synapse to the pacemaker group, providing important control mechanisms in the network Weaver and Hooper (2003); Mamiya and Nadim (2004). Furthermore, all the neurons in the network are strongly connected, so fluctuations in the activity of neurons that are not recorded can be detected in the activity of the LP and PD neurons.

From the continuous time series, we measured the four important discrete variables to describe the system (Figure 2): the duration of an entire cycle of the LP neuron *T*, the end of the LP burst *E*_*LP*_, the beginning of the PD burst *B*_*PD*_, and the end of the PD burst *E*_*PD*_. The reference for all these measurements is the time of first spike of the LP burst in each cycle. These variables define the onset and the termination of the recorded neurons bursts, and they have a clear behavioral significance: the LP and PD neurons are motoneurons and their burst determine the timing of activation and relaxation of two important pyloric muscles. We used

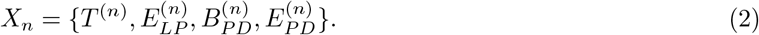

Hence,

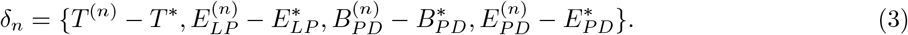

We calculated the stability of the system by computing the eigenvalues of **J** as a function of the strength of the synaptic connection between LP and PD neurons *g*_*syn*_. The synapse was artificially modified using the Dynamic Clamp hardware and software developed by Pinto et al. (2001). We varied the synaptic conductance *g*_*syn*_ from 0 to 100 nS. This parameter determines how much of the natural synapse is being reversed: 0 nS means intact synapse.

The distribution of the several eigenvalues obtained for one time series is shown in Figure 3. Since the time series are periodic, the module of the eigenvalues of the Jacobian must be smaller than 1. For the specific time series shown, there are two real and two complex eigenvalues, as shown by the four peaks in the histogram.

To address how our results depend on the size of the time series, we have first calculated the *λ* estimation with respect to the number of data points (Figure 4). In figure 4A we show the dependence of the probability of being unstable as a function of the number of data points. The value of *P*(∥*λ*_*max*_∥ ≥ 1|*N*) has an initial exponential decay on *N*. For sufficiently large *N, P*(∥*λ*_*max*_∥ ≥ 1|*N*) starts to saturate. In Fig. 4B we can see the mean curve and the 5% and 95% confidence level of the estimation of *λ* as a function of *N*. Its mean value converges quickly to an asymptotic value. The boundaries of the estimation of the maximal eigenvalue can also be exponentially reduced with the number of data samples. These results suggest that we can obtain reliable estimations even with small data sets.

**Figure 4:**
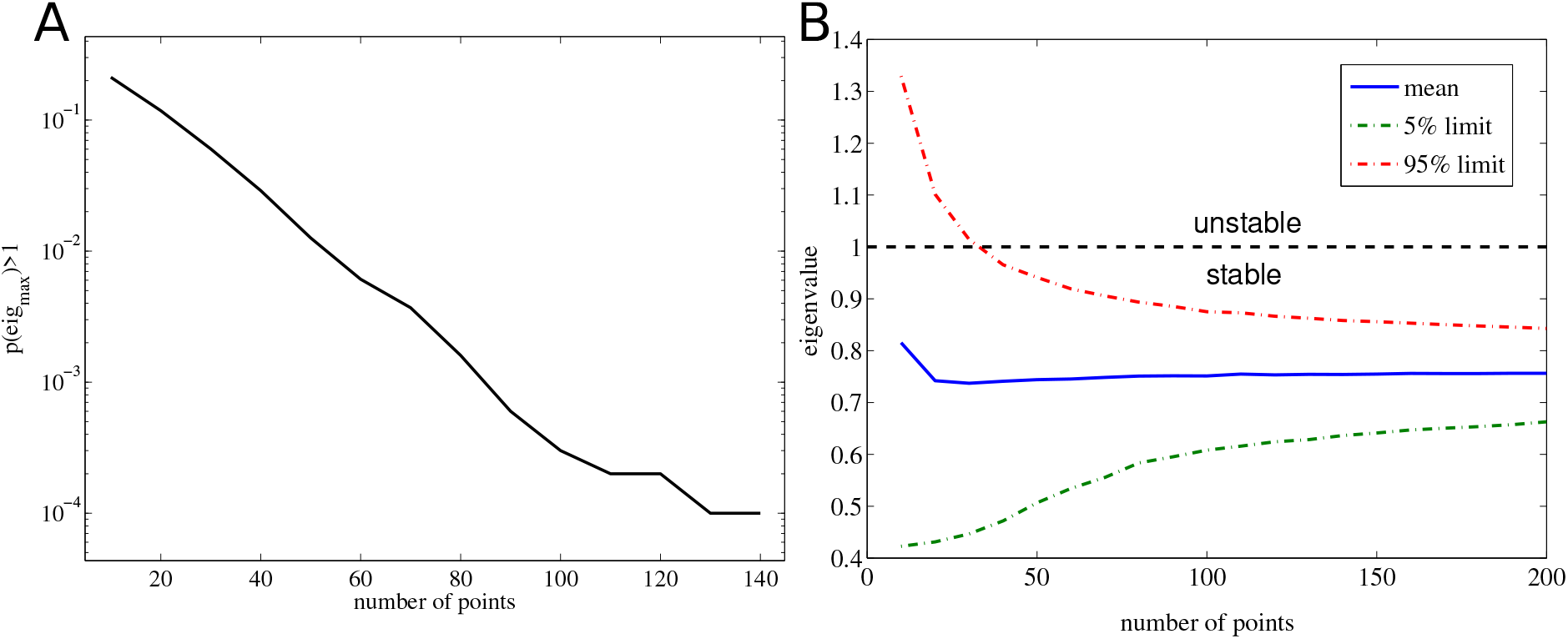
(A) Probability of having a eigenvalue larger than one by using *N* data points for the estimation. (B) Confidence interval as a function of the number of points considered for the eigenvalues calculation. The dash-dot lines show the 5% and the 95% confidence interval limits of the calculation.

Finally, we used the method to analyze the stability of the pyloric CPG with respect to a network modification of the synapse from the LP to the PD neuron. The eigenvalue that has the highest absolute value of its real part determines how close to a bifurcation the system is, and consequently, how stable it is. We show the results of these values in Figure 5. We observe that there is no significant modification of the system stability. Hence, we observe that the pyloric CPG stability is not affected by the modification of the LP to PD synapse. In Fig. 6 we show all the estimated eigenvalues for all the preparations and for all the synaptic changes. No *λ* value lies outside of the unit circle, so all of the CPGs are stable. No points were found outside of the [0, 1] range, showing that all the time series remained stable even under strong synaptic modifications. Finally, the upper limit of the 95% confidence interval of the eigenvalue was larger than 1 in 13 of the 55 data eigenvalues calculated. Even though in these cases we cannot rule out that the stability has changed, the lack of trend in the eigenvalues as a function of *g*_*syn*_, and the relatively large confidence intervals suggest that these cases are likely due to statistical fluctuations rather than an actual lack of stability.

**Figure 5:**
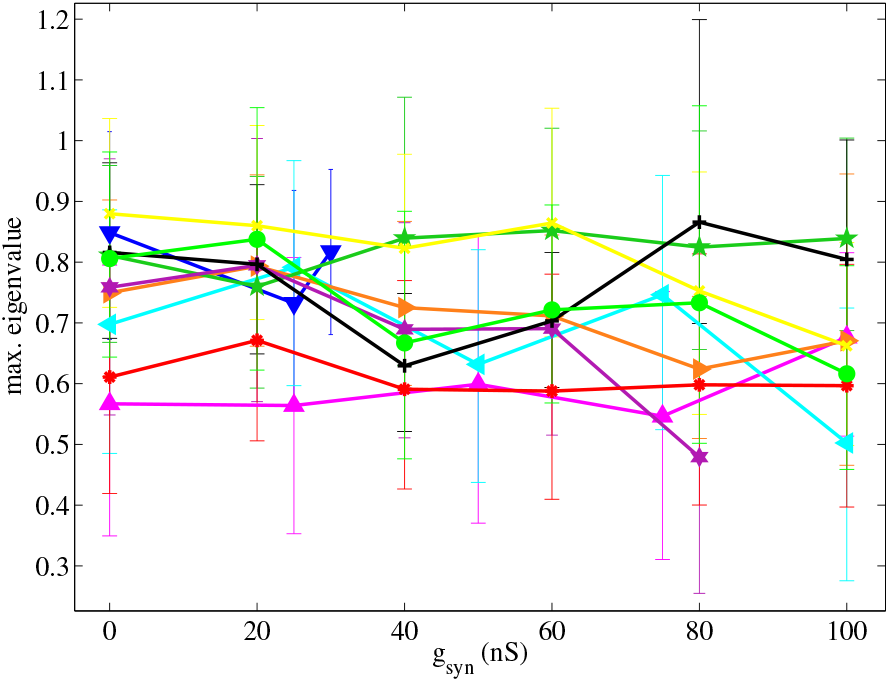
Maximum real part of the eigenvalues ∥*λ*_*max*_∥ as the synaptic conductance *g*_*syn*_ is changed for two preparations. Each curve represents a single preparation, and the error bars represent the 95% confidence interval. The maximum real part of the eigenvalues of the networks do not present a significant change when the synapse is reduced, showing that the stability of the oscillatory pattern is not modified by the synaptic modification.

**Figure 6:**
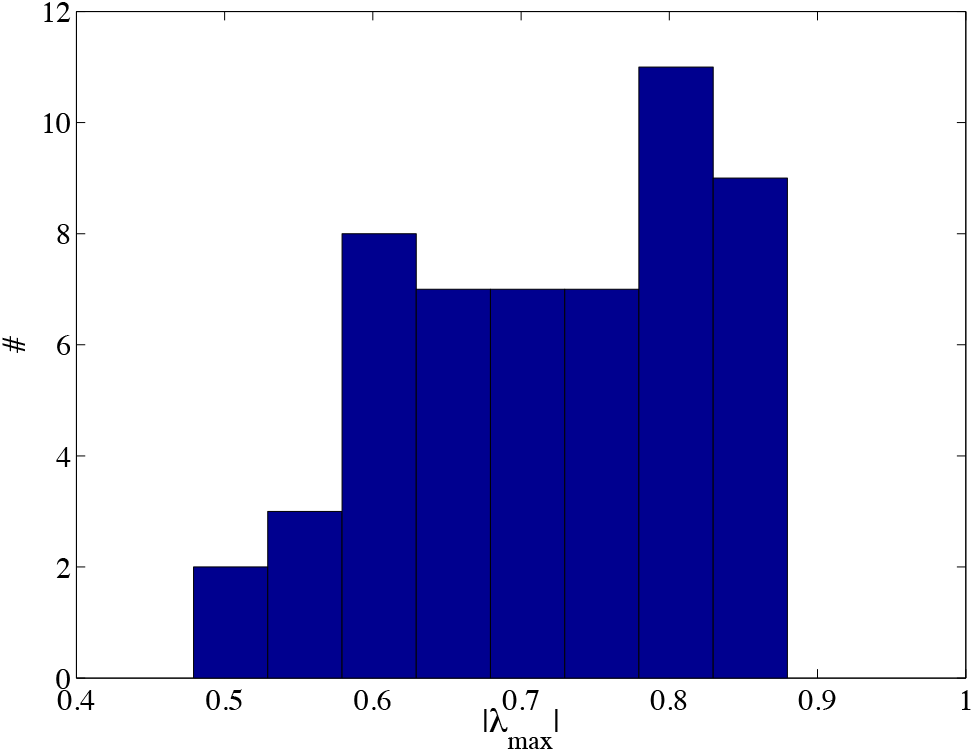
Histogram of the all the eigenvalues measured. We can see that all the experiments *λ* is in the interval [0.45, 0.9] showing that all the individuals presented a stable pattern even when the synaptic modification took place.

## Discussion

After decades of intensive study, there is an extensive literature on the fundamentals of the organization of CPGs Selverston et al. (2009); Selverston (1999); Marder and Calabrese (1996); Perkel and Mulloney (1974); Wang and Rinzel (1992); Skinner et al. (1994); Sharp et al. (1996); Rowat and Selverston (1997). In particular, much is known about the physiological properties of the crustacean pyloric CPG, both about the intrinsic neural properties and synaptic coupling. However, due to the complex nature of this system, the genesis of the rhythm is still not rigorously understood. Also, very little has been explored in terms of its dynamical stability. It is widely accepted that most stomatogastric neurons have intrinsic chaotic-like bursting activity when isolated Selverston et al. (2000); Carelli et al. (2005), and that the inhibitory synapses synchronize the neural activity, producing the stable rhythmic activity of the CPG Selverston et al. (2000); Reyes et al. (2015, 2008). The main points we address in this paper are the development of a novel method to infer the stability of CPGs based on its ongoing activity and to study how its stability is modified under synaptic modification via dynamic clamp.

In the present paper, we introduced a method to quantify the dynamical stability of an experimental dynamical system that mimics part of the pyloric CPG and is subject to intrinsic noise. We have shown how the noise in a dynamical system whose behavior was close to periodic could be used to gain information about the level of stability of this system. Similar methods to assess the eigenvalues of a linearized neural dynamics have been used in ECoG data, where the system is in a critical (not linearly stable) state with eigenvalues close to one Solovey et al. (2012, 2015) and simulated data with noise Escot and Sandubete (2023).

We also proposed a method that ascertains this stability with a given level of confidence. We used a bootstrap technique to quantify the confidence intervals of the estimation. In addition, it is interesting to note that, although the estimation of the eigenvalues to the data is straightforward, as far as we know, this is the first systematic quantification of the stability of rhythmic activity of CPGs.

We applied the method to data from the pyloric CPG, and tested its stability when the connectivity of the network was modified. We showed that the pyloric CPG stability in all preparations under analysis remained unaltered regardless of how much of the inhibitory synapse from the LP to the PD was canceled. This suggests that the pyloric CPG is densely connected with redundant synapses that keep the oscillatory behavior stable regardless of strong connectivity modifications. This could be one of the main principles under which these neural circuits are constructed (see O’Brien et al., 2025 for a computational approach to this question). If the CPG fails to deliver a rhythmic stable periodic neuronal firing, the proper coordination of the muscles cannot be achieved. Because of this redundancy, even removing a central synapse (such as the one from LP to PD) is insufficient to disrupt the normal pattern and the overall rhythm. Naturally, it must play another important role for the *in vivo* behavior, for example by finely tuning the LP/PD relative phase. The evidence we provide agrees with a number of studies showing that CPGs operate under dynamical invariants, i.e., certain phase-lags that scale with the rhythmic activity even for wide ranges of the main network frequency (Reyes et al., 2008; Elices et al., 2019; Reyes-Sanchez et al., 2023).

## Acknowledgments

This research was supported by NIH grant R01 NS050945, Fapesp grant 2022/16315-0, and CNpQ grants 430993/2016-1 and 408389/2024-9.

## Notes

### Competing Interest Statement

The authors have declared no competing interest.

### Summary of Updates

We have improved the two figures and included new references regarding the method used to quantify the stability. We also made small improvements throughout the text.

